# Thylakoid-integrated recombinant Hcf106 participates in the chloroplast Twin Arginine Transport (cpTat) system

**DOI:** 10.1101/382812

**Authors:** Qianqian Ma, Kristen Fite, Christopher Paul New, Carole Dabney-Smith

**Author notes:** Correspondence Carole Dabney-Smith, Miami University, 651 E. High St., Oxford OH, 45056. Current Address: Johns Hopkins University School of Medicine, Baltimore, MD. Current Address: Wright State University, Boonshoft School of Medicine, Dayton, OH.

## Abstract

The chloroplast Twin arginine transport (cpTat) system distinguishes itself as a protein transport pathway by translocating fully-folded proteins, using the proton-motive force (PMF) as the sole source of energy. The cpTat pathway is evolutionarily conserved with the Tat pathway found in the plasma membrane of many prokaryotes. The cpTat (E. coli) system uses three proteins, Tha4 (TatA), Hcf106 (TatB), and cpTatC (TatC), to form a transient translocase allowing the passage of precursor proteins. Briefly, cpTatC and Hcf106, with Tha4, form the initial receptor complex responsible for precursor protein recognition and binding in an energy-independent manner, while a separate pool of Tha4 assembles with the precursor-bound receptor complex in the presence the PMF. Analysis by blue-native polyacrylamide gel electrophoresis (BN-PAGE) shows that the receptor complex, in the absence of precursor, migrates near 700 kDa and contains cpTatC and Hcf106 with little Tha4 remaining after detergent solubilization. To investigate the role that Hcf106 may play in receptor complex oligomerization and/or stability, systematic cysteine substitutions were made in positions from the N-terminal transmembrane domain to the end of the predicted amphipathic helix of the protein. BN-PAGE analysis allowed us to identify the locations of amino acids in Hcf106 that were critical for interacting with cpTatC. Oxidative cross-linking allowed us to map interactions of the transmembrane domain and amphipathic helix region of Hcf106. In addition, we showed that *in vitro* expressed, integrated Hcf106 can interact with the precursor signal peptide domain and imported cpTatC, strongly suggesting that a subpopulation of the integrated Hcf106 is participating in competent cpTat complexes.

## 1 Introduction

The <u>T</u>win <u>A</u>rginine <u>T</u>ransport system (Tat) is one of two protein transport pathways that deliver proteins to the lumen of the plant thylakoid; a homologous Tat pathway is also found in a wide range of bacteria (Celedon and Cline, 2013; Berks, 2015; Cline, 2015). The Tat system is distinguished from other protein transport pathways, such as the well-characterized Secretory (Sec) system, by transporting fully-folded precursor proteins using only the proton-motive force (PMF) for energy (Cline and Mori, 2001; Braun et al., 2007; Gérard and Cline, 2007). Proteins transported by the Tat pathway are usually cytosolically synthesized as higher molecular weight precursors containing cleavable N-terminal signal peptides with an obligate twin arginine motif (RR) as is reflected in the name of the system. Tat systems can transport precursor proteins of different sizes ranging from 2 kDa to over 100 kDa, as well as substrates that form oligomers (Ma and Cline, 2010; Celedon and Cline, 2012). A recent stoichiometry study suggested that each individual Tat translocase can bind up to eight precursor proteins when fully saturated and is transport active when sufficient Tha4 is present (Celedon and Cline, 2012). However, mechanistic detail of how the receptor complex organizes to accommodate the transport of multiple precursor proteins in concert is an important question that needs to be addressed.

In thylakoid (as well as in *E. coli*) three membrane-bound protein components, Tha4 (TatA), Hcf106 (TatB), and cpTatC (TatC), are responsible for the twin arginine dependent translocation of precursor cargo (Cline and Mori, 2001; Celedon and Cline, 2013). Current models of the Tat system suggest a cyclical mechanism in which the receptor complex is comprised of cpTatC and Hcf106 tightly bound to each other with a loosely associated Tha4 that serves as the initial site of precursor recognition and binding, followed by assembly of additional Tha4 homo-oligomers, which are proposed to form the translocation pore (Bolhuis et al., 2001; Cline and Mori, 2001; Dabney-Smith et al., 2006; Dabney-Smith and Cline, 2009; Aldridge et al., 2012; Pal et al., 2013). Hcf106 is structurally similar to Tha4 in that both contain an amino terminal transmembrane domain (TMD), followed by a hinge region, an amphipathic α-helix (APH), and a loosely structured carboxyl terminus (C-tail). Recently, the structures of Tha4 and Hcf106 bacterial homologs, TatA and TatB, have been solved which agree with previous predictions (Hu et al., 2010; Zhang et al., 2014) and even more recently the structure to Tha4 from *Arabidopsis thaliana* (Pettersson et al., 2018). Despite the sequence similarity with Tha4, the two proteins are not interchangeable and thus appear to have distinct functions (Dabney-Smith et al., 2003).

Although the cpTat receptor complex has been well studied for its essential role in recognizing precursor proteins, few studies in plants have addressed the role of Hcf106 in this process, particularly, how each individual Hcf106 is organized in the multimeric receptor complex as compared to the bacterial homolog TatB (Bolhuis et al., 2001; Alami et al., 2003; Lee et al., 2006; Holzapfel et al., 2007; Rollauer et al., 2012; Behrendt and Bruser, 2014). The aim of the present work was to establish a method allowing the exploration Hcf106 organization using systematic cysteine substitution. As proof of principle, we have used this method to map Hcf106-Hcf106 interactions in thylakoid membranes.

Previous studies showed that *in vitro* translated Hcf106 can integrate into thylakoid in a manner presumably similar to endogenous Hcf106 and exists in a receptor complex with cpTatC (Gérard and Cline, 2006), which was capable of binding precursor proteins (Mori and Cline, 2002; Gérard and Cline, 2006). What remains unclear from the results of these studies is whether the integrated Hcf106 were simply members of the complex or directly participated in binding either precursor and/or cpTatC. Here, we have used cysteine scanning and disulfide bond formation to systematically map Hcf106 interactions through the TMD to the APH regions, which are known to be of great importance to the organization of the receptor complex. We observed that single cysteine-substituted Hcf106 protomers integrate into isolated thylakoid, that most variants are resistant to alkaline extraction. And that they localize in a 700 kDa complex by blue-native PAGE, suggesting that they are fully integrated into the membrane. Interaction sites of Hcf106-Hcf106 were obtained using copper (II)-1, 10-phenanthroline (CuP)-induced cross-linking which provided vital clues for the organization of Hcf106. Using double cysteine substitution in Hcf106, we could detect an Hcf106 oligomer as large as an octamer but could not distinguish if these oligomers were in the receptor complex or part of a separate pool of Hcf106. However, integrated Hcf106 was capable of interacting with transport competent precursor in a specific manner and with exogenous, imported cpTatC. From these data we conclude that integrated Hcf106 is capable of associating with and participating in the function of the cpTat translocase.

## 2 Materials and Methods

### 2.1 Preparation of chloroplasts and thylakoid membranes

Intact chloroplasts were prepared from 10-12 day-old pea seedlings (*Pisum sativum* L. cv. Laxton’s Progress 9 or Little Marvel) as described (Cline et al., 1993). Intact, isolated chloroplasts were suspended to 1 mg/ml chlorophyll in import buffer (IB, 50 mM HEPES-KOH, pH 8.0, 330 mM sorbitol) and kept on ice until used. Isolated thylakoid were obtained by osmotic lysis of intact chloroplasts. Briefly, intact chloroplast suspensions were pelleted for 5 min at 1000 xg, supernatant removed, and suspended at 1 mg/ml chlorophyll in lysis buffer (50 mM HEPES-KOH, pH 8.0, 10 mM MgCl_2_) with incubation on ice for 5 min. Following lysis, an equal volume of IB, 10 mM MgCl_2_ was added to the lysate, the thylakoid were then pelleted at 3200 xg for 8 min, and suspended at 1 mg/ml chlorophyll in IB, 10 mM MgCl_2_ (Aldridge et al., 2012). For single Cys interaction studies, thylakoid were suspended in 50 mM *N*-ethylmaleimide (NEM) in IB, 10 mM MgCl_2_ and incubated on ice for 10 min to prevent non-specific crosslinking from endogenous free sulfhydral groups. NEM-treated thylakoid were subsequently pelleted and washed with 3 volumes of IB, 10 mM MgCl_2_ before use.

### 2.2 Generation of Cysteine-substituted mature Hcf106 and C-tail truncation Hcf106_1-107_

Hcf106 with cysteine substitutions (Hcf106X_n_C; where amino acid X at position *n* was replaced by cysteine) were generated by QuikChange mutagenesis (Agilent Technologies) according to manufacturer’s instructions. The template used for mutagenesis was the coding sequence for mature Hcf106 (lacking the targeting peptide) in the plasmid pGEM-4Z. The coding sequence for Hcf106 begins with MASLFGVGAPEA…. Cloned constructs were verified by DNA sequencing on both strands at the Center for Bioinformatics and Functional Genomics at Miami University.

For the C-tail truncation of Hcf106_1-107_, internal stop codons were inserted via primer-based mutagenesis to generate the C-deletion of 69 amino acids. The size of the truncated Hcf106 is about 17kD by SDS-PAGE analysis.

### 2.3 Preparation of radiolabeled recombinant Hcf106 proteins

Radiolabeled Hcf106 variants were prepared by *in vitro* translation in a wheat germ extract from capped mRNA in the presence of [^3^H]leucine (Cline, 1986). The translation products were diluted with an equal volume of 60 mM leucine in 2× import buffer before use. Hcf106_1-107_ translation product was diluted further with 1× import buffer, 30 mM leucine equal to 1:3, 1:6, 1:12, or 1:60 of the initial translation before use.

### 2.4 *In vitro* integration assay

*In vitro* translated [^3^H]Hcf106X_n_C was integrated into NEM-pretreated thylakoid (100 μg chlorophyll equivalent) for 25 min at 25 °C. For Hcf106_1-107_, *in vitro* translated protein was directly integrated into isolated thylakoid. Reactions were terminated by transfer to 0°C and thylakoid were recovered by centrifugation at 3200 xg for 8 min. Recovered thylakoid were washed once with 2 volumes of IB, 10 mM MgCl_2_.

### 2.5 Alkaline extraction assay

Thylakoid with Hcf106 integrated were suspended to 1 mL with 0.2 M Na_2_CO_3_ or 0.1 M NaOl·l and incubated for 60 min on ice. Thylakoid were then recovered by centrifugation at 100,000 xg for 15 min. Pellets were suspended in 30 μl of 20 mM EDTA 1× IB and mixed with the same volume of 2× reducing sample solubilizing buffer (2× SSB (red); 100 mM Tris-HCl (pH 6.8), 0.2 M DTT, 5% SDS, and 30% glycerol). Samples were analyzed by SDS-PAGE and fluorography.

### 2.6 Blue-native gel electrophoresis

Hcf106 integrated thylakoid (~1 mg/ml chlorophyll) were dissolved in 2% digitonin with end-over-end mixing for 1 h at 4 °C and centrifuged at 100,000 xg for 30 min. Supernatant was mixed with 0.1 vol 10× BN-PAGE sample buffer (5% Serva G, 30% sucrose in 100 mM BisTris-HCl, 500 mM 6-amino-caproic acid, pH 7.0) as described by Cline and Mori (Cline and Mori, 2001). Gels were analyzed by fluorography or subjected to immunoblotting as described (Cline and Mori, 2001). Molecular markers used for blue native gels were dimeric and monomeric ferritin (880 kDa and 440 kDa, respectively) and bovine serum albumin (BSA) dimer (132 kDa).

### 2.7 Oxidative cross-linking by disulfide bond formation

Hcf106 integrated thylakoid were used for cross-linking reactions. 1 mM copper (II)-1, 10-phenanthroline (CuP from a 150 mM stock) was added as an oxidant to catalyze disulfide formation between proximal cysteine residues. The CuP stock solution (150 mM) contained 150 mM CuSO4 and 500 mM 1, 10-phenanthroline (Dabney-Smith et al., 2006). Cross-linking reactions were carried out for 5 min before stopping with 50 mM ethylmaleimide (NEM, from a 1 M stock in ethanol) and diluted two fold with 1× IB, 10 mM MgCl_2_. Thylakoids were recovered by centrifugation at 3200 xg for 8 min, the supernatants removed, and the pellet was suspended in 1× IB, 5 mM EDTA, 10 mM NEM and subjected to centrifugation and supernatant removal. Thylakoid pellets were suspended to ~1 mg/ml chlorophyll and divided into two, centrifuged at 3200 xg for 8 min, and the supernatants removed. The first half was suspended in 2× non-reducing sample solubilizing buffer (2× SSB (nr); 100 mM Tris-HCl (pH 6.8), 8 M urea, 5% SDS, and 30% glycerol) and the second half was suspended in 2× SSB (red). Samples then were analyzed by SDS-PAGE and fluorography. The intensities of the bands were quantified with ImageJ software (Schneider et al., 2012).

### 2.8 Disulfide cross-linking of imported cpTatC and integrated Hcf106

The precursor to cpTatC, pre-cpTatCaaaV270C (Kenneth Cline, University of Florida), was used as a source of exogenously integrated cpTatC according to published methods (Aldridge et al., 2014). Three native cysteines were substituted with alanine, hence the ‘aaa’ designation, and a single cysteine substitution was added at position V270. Radiolabeled *in vitro* translated pre-cpTatCaaaV270C was incubated with chloroplasts (0.33 mg/ml chlorophyll) and 5 mM Mg-ATP in IB with 100 μE/m^2^/s of white light in a 25°C water bath for 40 min. After import, intact chloroplasts were treated with thermolysin for 40 min at 4°C and isolated by centrifugation through a 35% Percoll (GE Healthcare) cushion in IB, 5 mM EDTA and washed with IB (Cline et al., 1993). Thylakoid membranes were prepared from isolated chloroplasts by osmotic lysis and centrifugation as described above and suspended in IB, 5 mM MgCl_2_ to ~1 mg/ml chlorophyll. *In vitro* translated, unlabeled Hcf106 was integrated into thylakoid as described above. Thylakoids were centrifuged at 3200 xg, 8 min to remove unincorporated Hcf106 and washed with IB, 10mM MgCl_2_. Samples were subjected to cross-linking as describe above.

### 2.9 Disulfide cross-linking of integrated Hcf106 and precursor

Radiolabeled, *in vitro* translated precursor tOE17-25C V-20F, containing an inserted cysteine 25 residues upstream from the signal peptide cleavage site, was incubated with thylakoids which had been pre-integrated with *in vitro* translated, unlabeled Hcf106 in IB, 10mM MgCl_2_ at 100 μE/m^2^/s of white light in a 25 °C water bath for 5 min. Samples were subjected to cross-linking as described above.

### 2.10 Supplemental Information

**Supplemental Figure S1**. Hcf106 with cysteine substitutions in the (A) N-terminus, (B) TMD, (C) hinge integrates into thylakoid and is resistant to alkaline extraction.

**Supplemental Figure S2**. Quantification of Hcf106 dimer formation in the TMD and APH.

**Supplemental Figure S3**. Most Hcf106 dimers disappear in the presence of the reducing agent, dithiothreitol (DTT).

## 3 Results

### 3.1 Single cysteine variants of Hcf106 integrate into thylakoid membranes and are resistant to alkaline extraction

Earlier work demonstrated that *in vitro* translated wild type Hcf106, which lacks native cysteines, was able to spontaneously integrate into isolated thylakoid and was resistant to alkaline extraction by either 0.2 M carbonate buffer (pH 9.5) or 0.1 M NaOH (pH 11.5) (Fincher et al., 2003). The NaOH treatment extracts proteins that are peripherally associated with membrane or are partially embedded into the membrane via a single transmembrane domain; whereas carbonate extraction is capable of stripping peripherally associated proteins and but not embedded proteins (Rolland et al., 2006).

We generated a series of single-cysteine substitutions from the predicted transmembrane domain to the end of the predicted amphipathic helix in Hcf106 (Figure 1) and investigated whether these cysteine substitutions, especially in the TMH, affected the integration of Hcf106 into the thylakoid membrane as compared to wild type.

**Figure 1.**
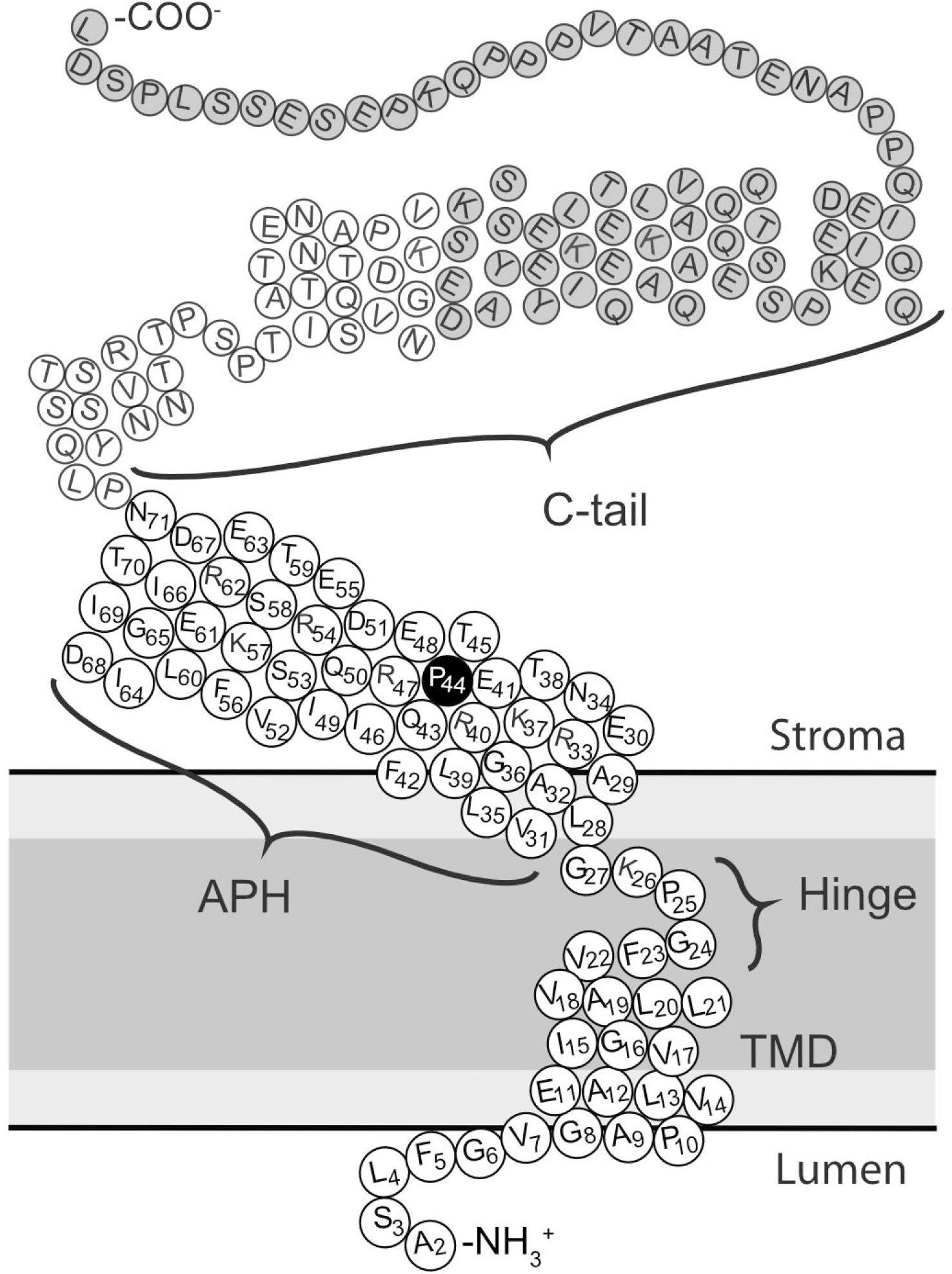
Topologic model of Hcf106 based on experimental and predictive structural features. Primary residue sequence of Hcf106 from *Pisum sativum.* Amino acids in regions under investigation in this study are numbered (2-71). Dark gray indicates the hydrophobic core (i.e. acyl chains) of the membrane, while the lighter gray indicates the lipid head groups.

Wild type Hcf106 is largely resistant to alkaline extraction (Fincher et al., 2003). Likewise, each of the Cys-substituted variants of Hcf106 tested were able to integrate into isolated thylakoid and were resistant to 0.2 M carbonate treatment. In addition, most of the variants were also resistant to 0.1 M NaOH extraction (Supplemental Figure S1); the exceptions are I15C, V17C, V18C, and L21C residues which are predicted to be in the hydrophobic core of the membrane. Thus, we concluded that the resistance of most single cysteine substituted Hcf106 to alkaline extraction indicates that the integration of recombinant Hcf106 into isolated thylakoids was not negatively affected.

### 3.2 Hcf106 Cys-variants can be detected in the 700 kDa receptor complex

Blue-native polyacrylamide gel electrophoresis (BN-PAGE) has shown that the cpTatC/Hcf106 receptor complex migrates as a band at ~700 kDa after solubilization by the detergent digitonin (Cline and Mori, 2001; Fincher et al., 2003). In addition, previous studies showed that recombinant, wild type, thylakoid integrated Hcf106 also migrated at 700 kDa, as well as a separate pool of smaller complexes of ~400 kDa to ~200 kDa depending upon the detergent to membrane ratio (Mori et al., 2001; Fincher et al., 2003). Most of the single cysteine variants of Hcf106 were integrated into isolated thylakoid; therefore, we examined whether the integrated Cys variants would form oligomers when solubilized in detergent as shown by BN-PAGE (Figure 2). We reasoned that if the Cys variants of Hcf106 could interact with cpTatC, then we would expect to find the Cys variants incorporated into a ~700 kDa complex when digitonin-solubilized thylakoid were analyzed by BN-PAGE.

**Figure 2.**
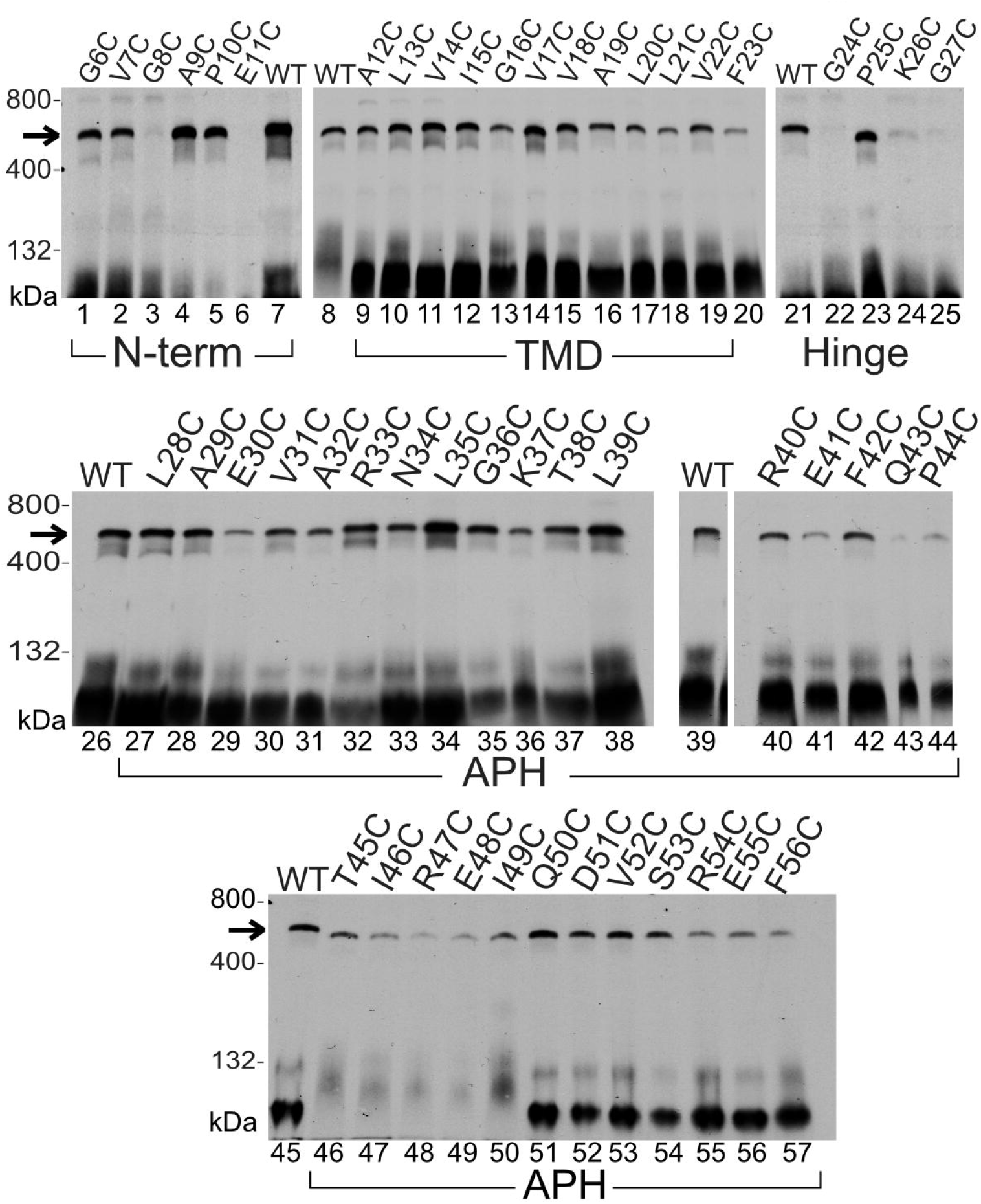
Blue-native gel analysis of the integration of recombinant Hcf106 cysteine variants. [^3^H]Hcf106 single cysteine variants containing complexes were analyzed by BN-PAGE and fluorography (Materials and Methods). Single Cys substitutions at multiple positions in Hcf106 are indicated across the top of the panels. The 700 kDa complex is indicated by an arrow. Wild-type Hcf106 was used as a control. Molecular mass markers are ferritin (880 and 440 kDa) and BSA (132 kDa). Gels are representative of at least three experiments.

We integrated the Hcf106 variants into isolated thylakoid and solubilized the membranes with digitonin followed with analysis by BN-PAGE. The ratio of detergent to thylakoid chlorophyll content was critical because previous studies demonstrated that an increase in the ratio of detergent to thylakoid resulted in the persistence of the 700 kDa receptor complex of cpTatC/Hcf106 complex, while the Hcf106 homo-oligomeric complexes between 400 kDa and 200 kDa were disrupted (Cline and Mori, 2001; Fincher et al., 2003). We were most interested in the presence or incorporation of Hcf106 in the 700 kDa receptor complex, so membrane solubilization was done at a ratio of 2% digitonin to 1 mg/ml chlorophyll to minimize the formation of the smaller homo-oligomeric complexes (Fincher et al., 2003). Bands seen at the bottom of the gel indicate the presence of smaller oligomers of Hcf106 (e.g., dimer to tetramer of ~60-120 kDa), but nothing in the 200-400 kDa range. Of the cysteines placed close to the N-terminus, such as G6C, V7C, G8C, A9C, P10C, and E11C, only G8C (Figure 2, lane 3) and E11C (Figure 2, lane 6) were unable to migrate in a 700 kDa complex but did integrate successfully into thylakoid (Supplementary Figure S1), suggesting that key contacts between Hcf106 and cpTatC were disrupted in those variants. Hcf106 variants with a Cys substitution in the transmembrane region such as A12C, L13C, V14C, I15C, G16C, V17C, V18C, A19C, L20C, L21C, and V22C (Figure 2, lanes 9-19), did migrate as a 700 kDa complex just like wild type Hcf106 (Figure 2, lanes 7-8, 21, 26, 39, 45) suggesting incorporation into the receptor complex. Most of the variants in the hinge region, i.e., F23C, G24C, K26C, or G27C (Figure 2, lanes 20, 22, 24, and 25) failed to incorporate into the receptor complex but were able to integrate into thylakoid (Supplemental Figure S1). P25C was able to incorporate into the receptor complex (Figure 2, lane 23). Cys variants in the APH region of Hcf106 showed a similar pattern to wild type when analyzed by BN-PAGE in that all single cysteine substitutions in this region did not abolish incorporation into the 700 kDa complex (Figure 2, lanes 26-57). However, certain cysteine substitutions, for example, E30C, A32C, K37C, E41C, Q43C, and P44C (Figure 2, lanes 29, 31, 36, 41, 43, and 44), consistently showed a lower intensity at the 700 kDa band, suggesting that cysteines in this region of the APH may negatively affect Hcf106 interaction with cpTatC (i.e., incorporation into the 700 kDa complex), but did not affect the integration and membrane stability of the variant (Supplemental Figure S1).

### 3.3 C-terminal of Hcf106 is dispensable for cpTatC-Hcf106 receptor complex formation

Hcf106 contains a loosely-structured C-tail that was shown to not be required for receptor complex formation in the *E. coli* homolog, TatB (Maldonado et al., 2011). If the truncated Hcf106 could interact with cpTatC, we reasoned that truncated protein could be used to demonstrate that the integrated Hcf106 is, in fact, incorporating into receptor complexes with endogenous cpTatC (Figure 3). We incubated increasing amounts of [^3^H]Hcf106_1-107_, lacking 69 amino acids from the C terminus, with isolated thylakoid and analyzed the membranes by BN-PAGE. As the concentration of *in vitro* translated [^3^H]Hcf106_1-107_ increased, two lower bands of ~600 kDa (Figure 3, lanes 2-4) and 500 kDa (Figure 3, lanes 3-4) appeared, suggesting that *in vitro* integrated [^3^H]Hcf106 was competing with the endogenous Hcf106 for binding to endogenous cpTatC. The smaller complexes also contain cpTatC (Figure 3, lanes 9-12), indicating the C-tail does not play a critical role in receptor complex formation. Immunodetection of Hcf106 using the same thylakoid samples demonstrated that full length Hcf106 also migrated into smaller complexes due to the presence of the truncated variant. With these data, the insensitivity to alkaline extraction, and the incorporation into a 700 kDa complex, we conclude that exogenously added Hcf106 is properly inserted into the thylakoid membrane, allowing us to use these Cys-substituted variants to probe the organization of Hcf106 in the receptor complex.

**Figure 3.**
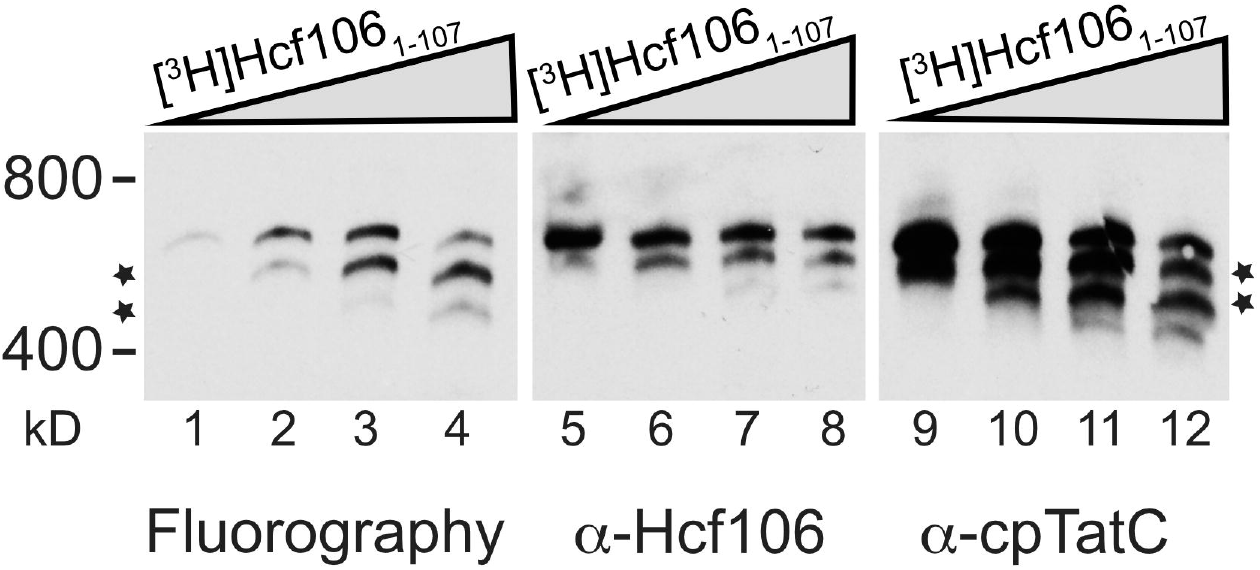
Truncated Hcf106_1-107_ assembles into a complex with endogenous cpTatC. An increasing concentration of Hcf106_1-107_ was integrated into thylakoid and subjected to digitonin solubilization, BN-PAGE, and analysis by fluorography (left panel) as in Figure 2. To detect whether this truncated form can generate a different size of the receptor complex, samples were also subjected to immunoblotting with anti-cpTatC (middle panel), anti-Hcf106 (right panel) antibodies. The amount of Hcf106_1-107_ added to thylakoid corresponds to dilutions of the *in vitro* translation reaction (i.e., 1:60, 1:12, 1:6, and 1:3). Gels are representative of at least three separate experiments. As Hcf106_1-107_ increases, there is a concomitant increase in a smaller complex also containing cpTatC (asterisks).

### 3.4 The transmembrane domain and amphipathic helix regions of Hcf106 form self-contacts

To characterize the organization of Hcf106-containing complexes, we looked at the organization of Hcf106 by studying interactions between neighboring Hcf106 proteins. We reasoned that interactions between Hcf106 proteins would indicate the organization of Hcf106 in the receptor complex by identifying sites specific for self-interactions as well as provide insight into the organization of the separate pool of Hcf106. We took a cysteine scanning approach, which allowed us to map interactions between neighboring single cysteine substituted Hcf106 proteins or other cpTat components by formation of disulfide bonds between cysteines within ~5 Å of each. Hcf106 proteins containing single cysteine substitutions in the TMD or APH were integrated into isolated thylakoid. In the presence of an oxidant such as copper (II)-1,10-phenanthroline (CuP), free cysteine sulfhydrals in close proximity will form stable disulfide bonds, which cause a mobility shift from ~28 kDa to ~56 kDa of the crosslinked proteins when analyzed by SDS-PAGE.

Residues close to the N-terminus of Hcf106 were in close proximity to the same residue of a neighboring Hcf106 (Figure 4, lanes 1-5) showing a significant amount of dimer formation, demonstrating that these amino acids are sufficiently close to form a disulfide bond or that this region is very flexible. On the other hand, when the Cys was placed in the TMD, i.e., E11C-L21C (Figure 4, lanes 6-16), dimer formation demonstrated a position-specific interaction, indicating a regular face of interaction in this portion of the transmembrane domain. As the cysteine substitution moved out of the TMD through the hinge and into the APH, i.e., V22C, F23C, and K26C (Figure 4, lanes 17, 18, 21) formed a dimer, whereas G24C, P25C, and G27C (Figure 4, lanes 19, 20, 22) did not.

**Figure 4.**
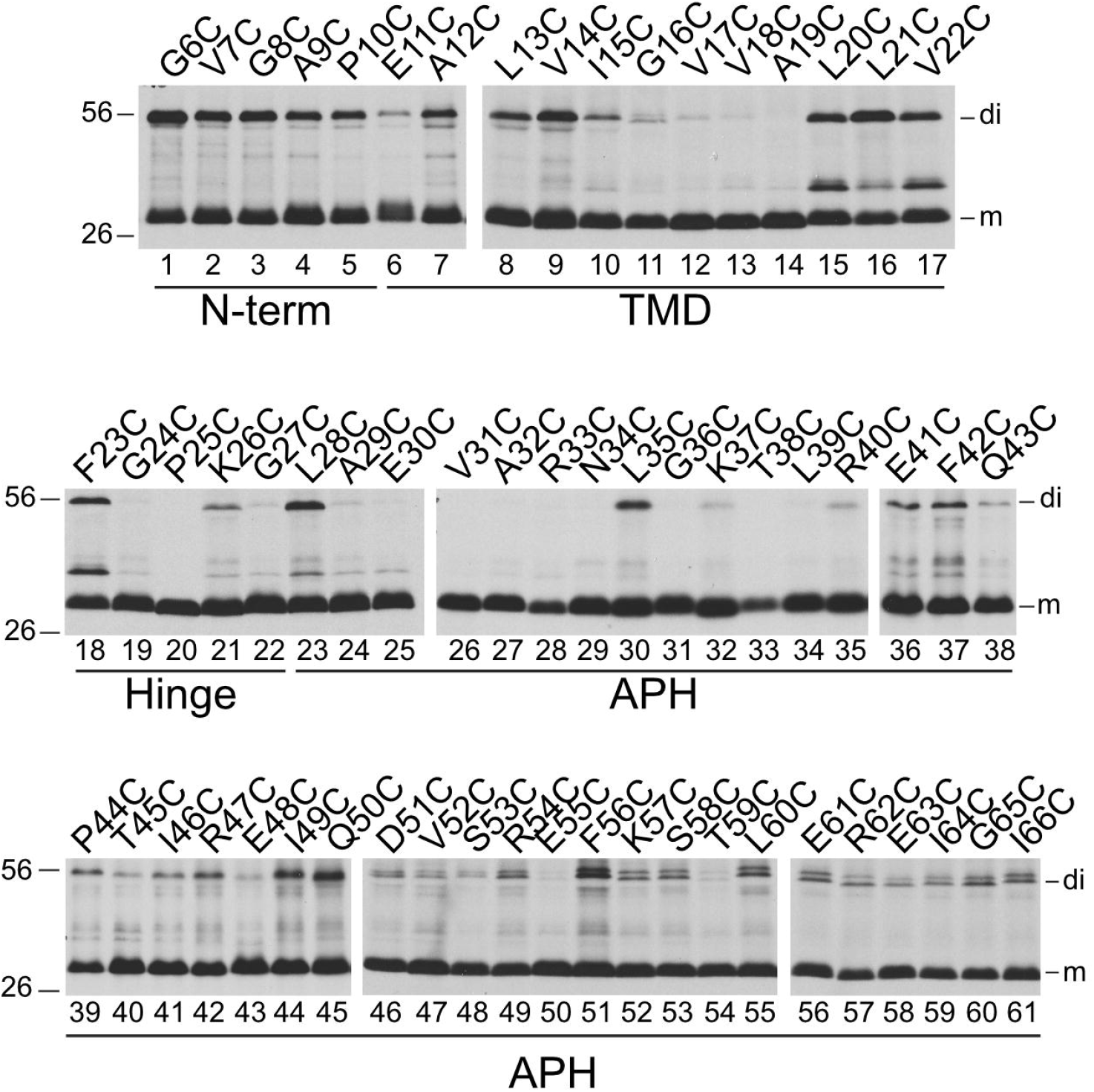
Hcf106 forms a dimer when cysteine substitutions are in N-terminus, the central TMD, and the APH regions. *In vitro* translated Hcf106 variants were integrated into NEM pre-treated thylakoid membranes and subjected to oxidizing conditions (1 mM CuP) as described (Materials and Methods). Samples were resolved by SDS-PAGE and protein bands were visualized by fluorography. The amino acid position of each cysteine substitution is shown above each panel. Hcf106 monomer (m) and dimer (di) forms are indicated at the right of the panels. Gels are representative of at least three separate experiments.

The amphipathic helix of Hcf106 showed two different types of interaction. The N-terminal proximal portion of the APH, i.e., L28C-P44 (Figure 4, lanes 23-39), showed no interaction overall, with the exceptions of L28C, L35C, E41C, and F42C (Figure 4, lanes 23, 30, 36, 37), compared to the C-terminal proximal portion of the APH, T45C-G65C (Figure 4, lanes 40-60), which demonstrated a stronger and position-specific cross-linking. For example, dimer was detectable for all Cys-substitutions in this segment, but dimers formed with cysteines at I49C or Q50C were more intense (Figure 4, compare lanes 44-45 with lanes 36-43) based on equal chlorophyll loading. The relative proportion of dimer to total protein was quantified by densitometric analysis (Supplemental Figure S3). The presence of a band at 56 kDa was dependent upon disulfide formation because treatment of samples with the reducing agent, dithiothreitol (DTT), effectively depleted the dimer species. Interestingly, L21C and V22C were not reduced to the monomeric form (Supplemental Figure S3).

The results of the specific interactions on the Hcf106 TMD or APH are plotted on helical wheels (Figure 5). For example, a helical wheel projection of the TMD with the hinge region (residues A12C-G27C) emphasizes that Hcf106 self-interactions occur along no particular face (Figure 5A). The APH domain (~40 amino acids) is too large to be clearly evaluated with one helical wheel, so we generated wheels for the N-terminal half (i.e., L28-P44; (Figure 5B) and the C-terminal half (T45-G65; Figure 5C). The proline at position 44 was arbitrarily determined as the halfway point because it would serve to break the helix. The helical wheel of the N-terminal portion of the APH shows a preferred face for Hcf106 self-interaction because L28, L35 and F42 fall along the hydrophobic face of the helix (Figure 5B), although we do see interactions at E41C and P44C, which do not fall on the same face of the predicted helix. However, we envision the APH could be very mobile or dynamic in the membrane, which may explain the interactions at E41C and P44C. Other residues, e.g., A29C-T38C, showed no self-interactions indicating that these faces of the helix may be buried in the membrane, interacting with other cpTat components, or positioned such that they are not near each other on neighboring helices. The helical wheel of the C-terminal portion of the APH demonstrated that roughly three of four faces can interact with neighboring Hcf106 (Figure 5C), indicating that this portion of the helix might be flexible or that it could emanate from the membrane similarly to what was seen with Tha4 (Aldridge et al., 2012).

**Figure 5.**
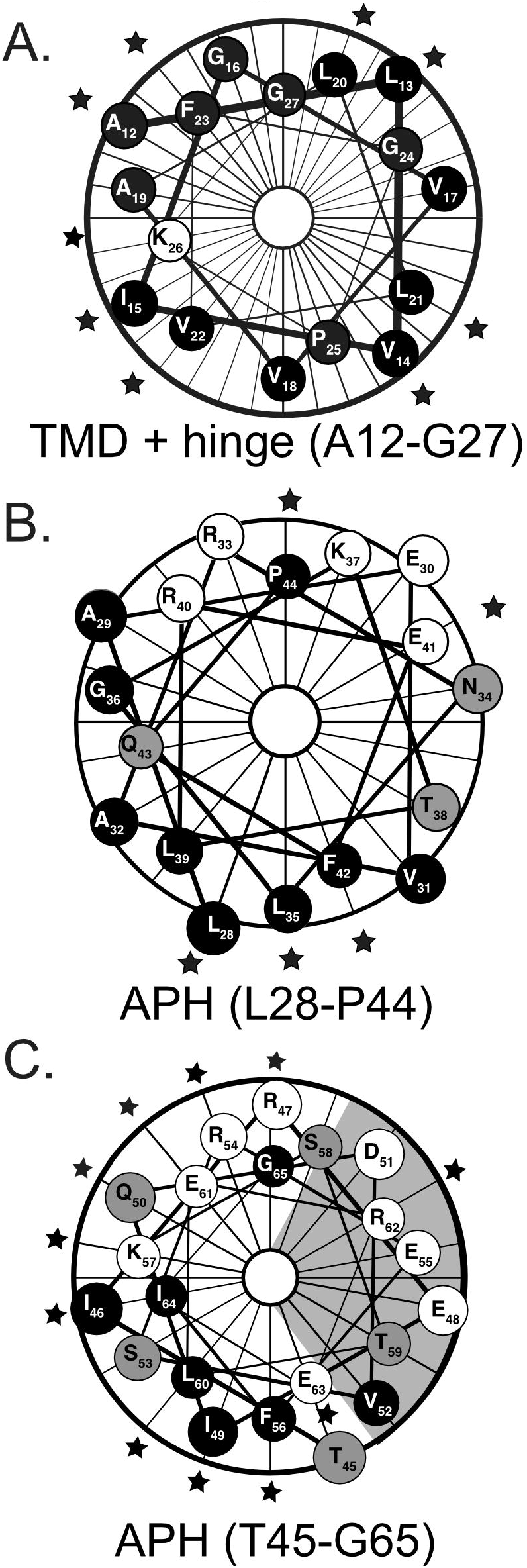
Helical wheel projection of the TMD and APH regions of Hcf106 reveals periodic interactions. Helical wheel projections were generated using the Protean module of DNAStar (Lasergene, Madison WI). (A) The TMD and Hinge of Hcf106, residues A12C-G27C; (B) the N-terminal proximal portion of the APH of Hcf106, residues L28C-P44C; and (C) the C-terminal proximal portion of the APH of Hcf106, residues T45C-G65C. The shading represents the hydrophobicity of the amino acid, hydrophilic amino acids are more lightly shaded. Stars indicate the presence of a dimer that is >10% of total (see Supplemental Figure S2).

### 3.5 Double-cysteine substitutions suggest a high-ordered Hcf106 complex

Single cysteine substitutions can only detect dimers. To investigate the formation of higher-ordered oligomers, we constructed double cysteine substituted Hcf106 to detect an ability to form higher ordered protomers. We substituted two cysteines in the transmembrane helix of Hcf106, e.g., l13CL21C and V14CL20C, and the resulting proteins were integrated into thylakoid followed by oxidative crosslinking. Initially, when compared to the single cysteine variant L21C, the double Cys-substituted proteins L13CL21C and V14CL20C showed roughly half the integration into thylakoid as L21C (Figure 6A, compare lanes 7, 10 to 9, 12), while double cysteine variants in the APH, e.g., I49CE61C and R54CL60C (Figure 6A, compare lanes 8, 11 to lanes 9, 12), were comparable to the amount of L21C integrated. Decreased integration of the TMD double Cys variants was likely due to the introduction of two cysteines into the helix, decreasing the hydrophobicity of the TMD. Unexpectedly, the cross-linking of these four cysteine variants did not show obvious Hcf106 multimers (Figure 6A, lanes 1-2, 4-5). Furthermore, when the V14CL20C variant was compared to its single cysteine parent, V14C, the double Cys variant also showed about 30% less integration into thylakoids (Figure 6B, lanes 1-2). However, when one cysteine was placed in the N terminus and the other in the TMD, e.g., G6CV14C, we observed Hcf106 multimers as high as octamers when analyzed by 10-20% acrylamide gradient gel PAGE (Figure 6, lanes 34).

**Figure 6.**
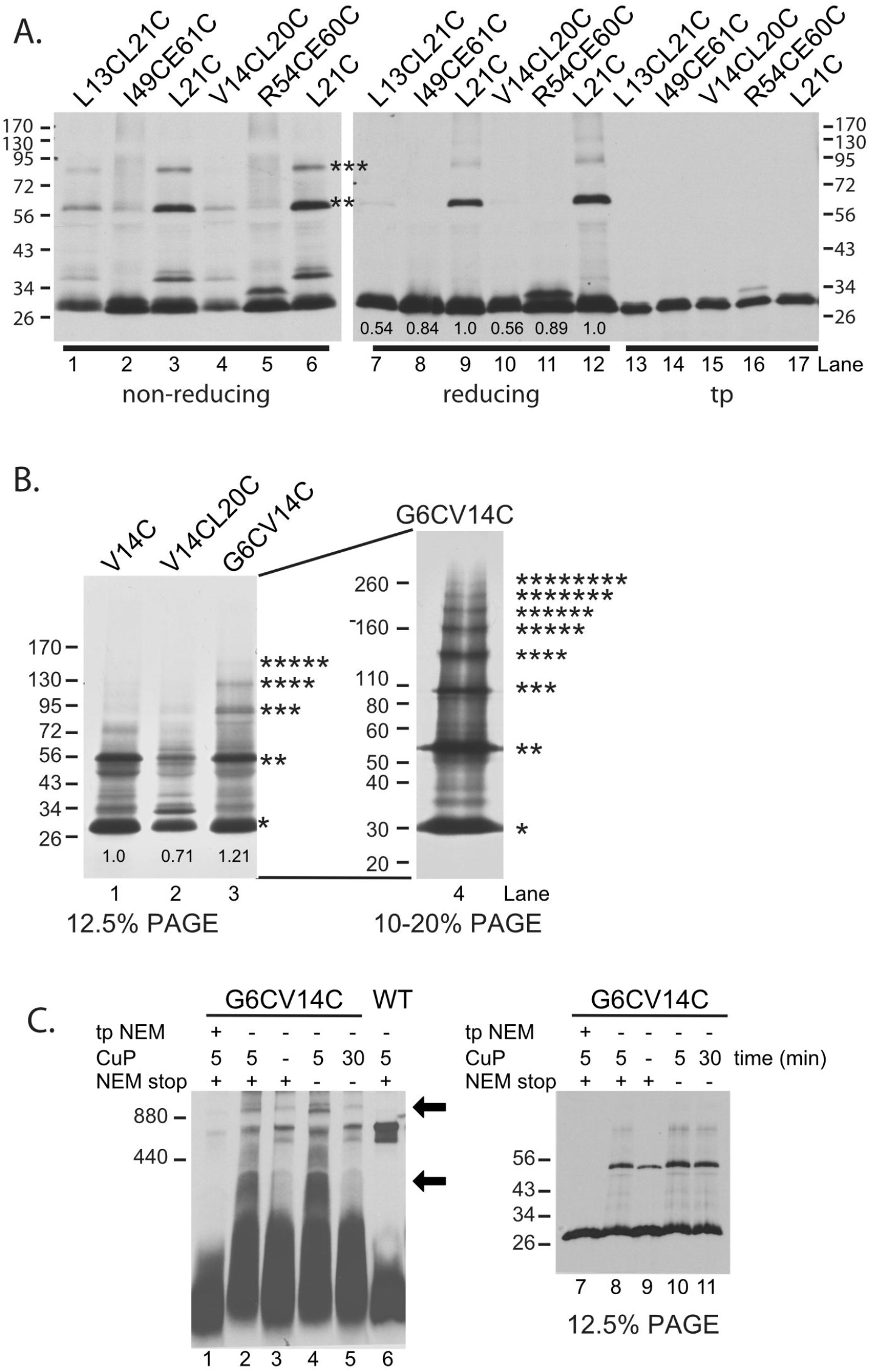
Double cysteine substitutions demonstrate a higher order Hcf106 complex. (A) Lanes 1-6 show the cross-linking of double cysteine variants L13CL21C, I49CE61C v14CL20C and R54CL60C and the single Cys variant L21C under nonreducing conditions. Lanes 7-12 show the same samples under reducing conditions. The amount of protein integrated relative to the parent single Cys variant, L21C, is shown below lanes 7-12. Translation products diluted 1:40 are shown in lanes 13-17. Molecular weight markers are indicated on the right and left. (B) Crosslinking with the G6CV14C double cysteine variant was compared to the single Cys mutant V14C and double mutant V14CL20C (lanes 1-3). G6CV14C showed higher ordered Hcf106 complexes up to octamers when analyzed by 10-20% SDS-PAGE (lane 4). (C) Crosslinked G6CV14C double Cys variant has reduced capability to incorporate into the receptor complex. Lane 2 is the standard cross-linking condition with 5 min incubation of CuP and the reaction was quenched with 50mM NEM. Lane 1: translation product was pre-treated with NEM and subjected to standard cross-linking. Lanes 3-6: cross-linking under variable time or with/without NEM quenching. Lane 6: WT Hcf106 under standard cross-linking conditions. The same samples from lanes 1-5 were also subjected to nondenaturing SDS-PAGE (lane 7-11). Asterisks indicate the number of Hcf106 monomers in the oligomer as determined by size. Arrows indicate additional bands of cross-linked G6CV14C. The amount of protein integrated relative to the parent single Cys variants, V14C, is shown below the lanes. Gels in both panels represent at least three separate experiments.

To further investigate if the cross-linked Hcf106 multimers are part of the 700 kDa complex, we subjected the cross-linked G6CV14C double Cys variant to BN-PAGE (Figure 6C). Different cross-linking conditions were used including pre-treatment of the translation product with NEM to block the free cysteine (Figure 6, lane 1), the presence or absence of CuP (Figure 6, lanes 2 and 3), cross-linking without quenching (Figure 6, lane 4), and prolonged cross-linking (30 min) (Figure 6, lane 5). Samples were also subjected to non-reducing SDS-PAGE to monitor the status of cross-linking (Figure 6, lane 7-11). On BN-PAGE, we observed a significant amount of 700 kDa complex when the double cysteine variant was cross-linked (Figure 6, lane 2-5). Additional bands were also observed (above 800 kDa and <400 kDa, indicated with arrows), especially when cross-linking in the absence of NEM quenching (Figure 6, lane 4). The bands above the 800 kDa could be non-specific interactions between the 700 kDa complex and other species. It could also be due to the different mobility of the complex caused by crosslinking induced conformational change. While we interpret the <400 kDa band to be indicative of the G6CV14C variant in the separate Hcf106 pool, which may also form higher oligomers. Overall, these results indicate Hcf106 has a strong tendency to form oligomers and that Hcf106 self-oligomerization might be present in both the receptor complex and the free pool of Hcf106.

### 3.6 Integrated Hcf106 form contacts with both imported cpTatC and precursor proteins

To validate whether integrated Hcf106 can participate in the transport process directly, we looked for contacts between integrated Hcf106 and imported cpTatC or cpTat pathway precursor proteins. Previous studies found that endogenous Hcf106 can be photo-crosslinked with the signal peptide of precursor proteins when part of a functional receptor complex with cpTatC (Gérard and Cline, 2006), and that integrated, recombinant Hcf106 localizes to a complex that could bind precursor (Cline and Mori, 2001).

Earlier work demonstrated that replacing the original cpTatC transit peptide with the transit peptide of the precursor to the small subunit of RuBisCO resulted in higher efficiency of pre-cpTatC import, mature cpTatC localization to thylakoid that could be detected in direct contact with both the RR proximal region on a Tat pathway precursor signal peptide and Tha4 (Aldridge et al., 2012; Ma and Cline, 2013; Aldridge et al., 2014). To confirm Cys-substituted Hcf106 participation in the receptor complex, we selected five Hcf106 Cys variants from the transmembrane domain and looked to see if they interact with imported cpTatCV270C. We initially chose transmembrane locations for cysteine substitution because recently the cpTatC bacterial homolog, TatC, was found to form crosslinks with the TatB (Hcf106) transmembrane domain via TatC TM5 (Kneuper et al., 2012; Rollauer et al., 2012). Imported cpTatC270C was previously used to map Tha4 binding (Ma and Cline, 2013). We observed [^3^H]cpTatC at an apparent molecular weight of ~28 kDa, which matches the apparent molecular weight for endogenous cpTatC. Samples with integrated Hcf106, containing Cys-substitutions in the hydrophobic core of the Hcf106 transmembrane domain, e.g., A12C, L13C and V14C, formed interactions with cpTatCV270C, showing a ~56 kDa adduct that is not found in the control lane (Figure 7A). For the control lane, no Hcf106 was integrated into the thylakoid. In contrast, L20C, which is closer to the stromal side of the membrane, showed a similar result as the control lane. The data here demonstrate that integrated Hcf106 is in close contact with cpTatC.

**Figure 7.**
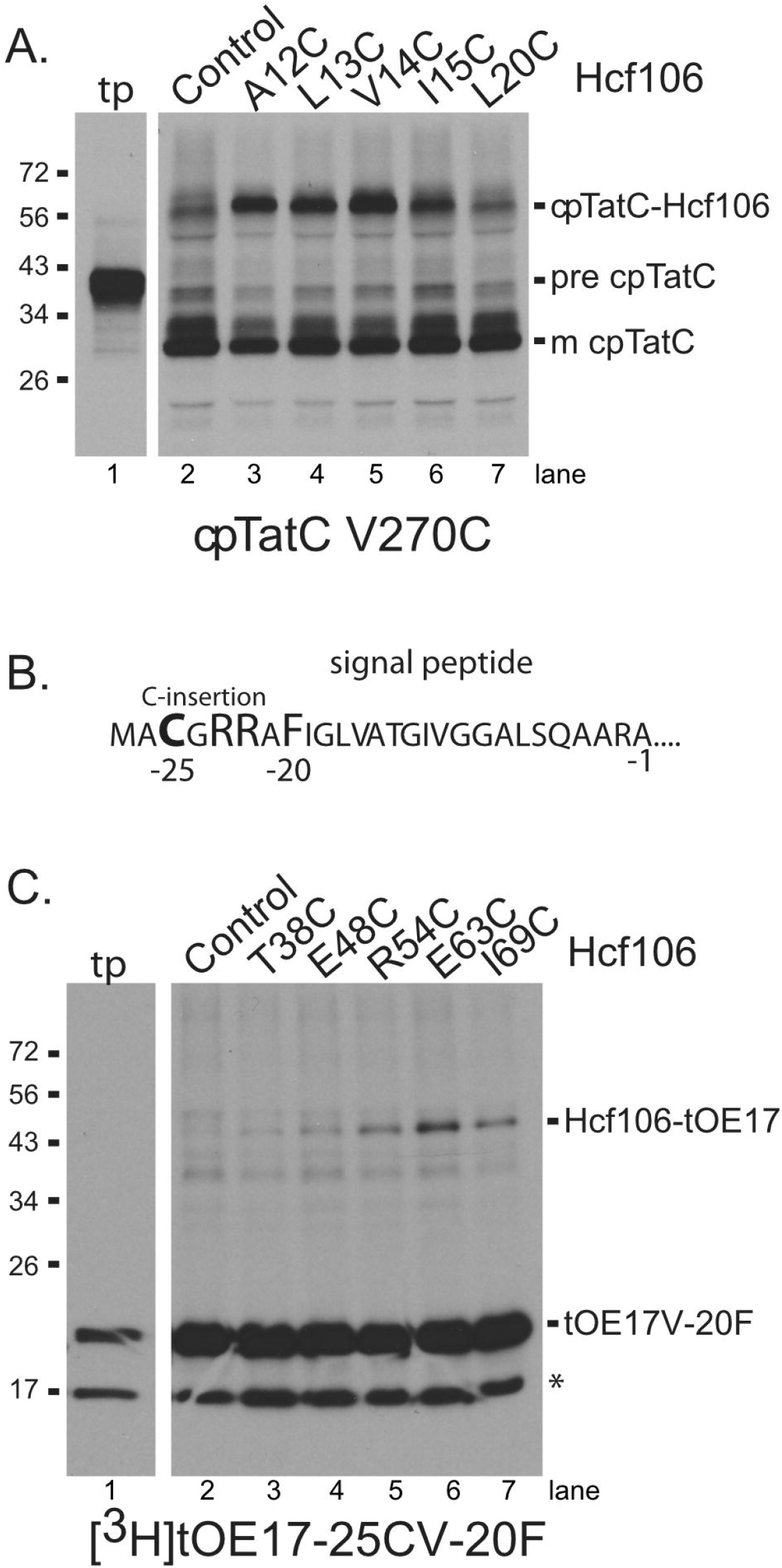
Integrated Hcf106 interacts directly with imported cpTatC and the signal peptide of the precursor. (A) *In vitro* translated [^3^H]cpTatCaaaV270C was imported into intact chloroplasts. Intact chloroplasts were isolated over a 35% Percoll cushion (Materials and Methods) and recovered thylakoid were used for integration of unlabeled Hcf106 Cys variants, as indicated across the top of the panels, or the same volume of IB, 10 mM MgCl_2_ as control. Hcf106 TMD shows cross-linking with cpTatC TM5 in a ~56 kDa band that migrates higher than the nonspecific bands in the control lane. Translation products (tp) diluted 1:40 are shown. (B) Residue sequence of the signal peptide of tOE17-25CV-20F showing the -25C substitution, the RR motif, and the V-20F substitution. (C) *In vitro* translated unlabeled Hcf106 Cys variants, as indicated across the top of the panel, were integrated into isolated thylakoids. Thylakoid were then washed to remove unintegrated protein (Materials and Methods) and incubated with either precursor, [^3^H]tOE17-25CV-20F, or the same volume of iB, 10 mM MgCl_2_ as control. Gels in both panels are representative of at least three separate experiments.

Alternatively, to confirm that integrated, recombinant, Cys-substituted Hcf106 was able to bind precursor, we integrated wild type Hcf106 or various Cys-substituted Hcf106 into thylakoid and used these membranes in precursor binding assays. The precursor, tOE17-25C/V-20F, was shown to bind tightly to the first cpTatC stromal loop (Gérard and Cline, 2007) and was used previously cpTatC crosslinking assays (Ma and Cline, 2010). It is a modified precursor of the 17 kDa subunit of the oxygen evolving complex, containing a truncated signal peptide (tOE17), a cysteine inserted on the N-terminal side of the twin arginine motif, 25 amino acids from the signal peptide cleavage site (-25C), and a phenylalanine substituted for the valine at position -20 from the signal peptide cleavage site (V-20F) [Figure 7, see reference (Ma and Cline, 2010). We subjected tOE17-25C/V-20F to cross-linking with the Hcf106 C-terminal APH region as it interacts with cpTatC stromal loop 1 (Figure 7C). This loop of cpTatC was also identified as interacting with tOE17-25C/V-20F. tOE17-25C/V-20F showed an interaction with Hcf106T38C, E48C, R54C, E63C, and I69C by the presence of a higher molecular weight adduct when analyzed by non-reducing SDS-PAGE. The strongest interactions involved Hcf106R54C, E63C, and I69C (Figure 7C, lanes 3-7), while wild type Hcf106, lacking cysteine, was not able to generate an adduct (Figure 7C, lane 2). Taken together these data suggest that Hcf106 variants are incorporating into receptor complexes and can play a functional role in precursor binding.

## 4 Discussion

Recently, structural insights of the individual prokaryotic Tat components were revealed which lay a solid foundation for deciphering the mechanism of the Tat system (Hu et al., 2010; Rollauer et al., 2012; Ramasamy et al., 2013; Zhang et al., 2014). Additionally Blümmel *et al.* used photo-crosslinking to elucidate the architecture of TatBC oligomers during the initial assembly step, but due to the nature of the technique could only provide specific residue location for one binding partner (Blummel et al., 2015). Here, cysteine scanning by disulfide bond formation was used to provide a precise map of interactions between the two homologous proteins in thylakoid. Previous work using isolated thylakoids demonstrated a substrate-gated docking of Tha4 into the cpTatC cavity initiating translocase assembly (Aldridge et al., 2014) and challenging the receptor complex model from *E.coli* in which TatB forms a ring-like structure in the center with TatC occupying the peripheral positions (Maurer et al., 2010; Cline, 2015). However, how Hcf106-cpTatC are arranged to accommodate Tha4 docking and whether there are organizational differences between TatBC in *E. coli* and Hcf106-cpTatC in thylakoid have not been determined yet due to a lack of methodology to study the role of Hcf106 in isolate membranes.

Despite sharing significant sequence similarity, Tha4 and Hcf106 have distinct roles in protein transport (Cline and Mori, 2001; Dabney-Smith et al., 2003). Unlike Tha4, Hcf106 is typically found tightly complexed with cpTatC. Therefore, the use of the α-Hcf106 antibody is not viable as a sequestrant for native Hcf106 as it was for Tha4 (Dabney-Smith et al., 2003) due to the tight association of Hcf106 with cpTatC. Previous experiments showed that integration of *in vitro* translated [^3^H]Hcf106 into thylakoid assembled into a 700 kDa receptor complex as analyzed by BN-PAGE (Fincher et al., 2003), setting the stage for a possible involvement of exogenously Hcf106 in the receptor complex. Here we further characterize integration of *in vitro* expressed Hcf106 using cysteine substitutions and disulfide bond formation to demonstrate self-interactions and interactions with imported cpTatC and precursor. Introduction of cysteine substitutions into Hcf106 largely did not affect integration into the membrane or presence in the 700 kDa complex. Exceptions include Cys-substitutions in areas such as the TMD hydrophobic core, which lowered integration overall, likely due to a decrease in helix hydrophobicity, and in the hinge region, which may also be buried or involved in contacts with cpTatC. However, most of the Hcf106 Cys-substitutions tested do integrate into thylakoid allowing the study of the organization of the protein in a native membrane. In *E. coli,* in the absence of TatC, TatB formed a ladder of bands of about 100 kDa to over 880 kDa, suggesting that TatB has oligomeric properties on its own when it is not associated with TatC (Behrendt et al., 2007; Cleon et al., 2015). In the present study, we see ladders of full length and truncated Hcf106, which may be indicative of the separate pool of Hcf106; however, we also see formation of ladders of cpTatC that correspond in a linear manner to the integration of the truncated version of Hcf106. We conclude, therefore, that a substantial fraction of the incorporated, truncated Hcf106 is a part of the receptor complex with endogenous cpTatC.

We also identified residues in Hcf106 that are important to receptor complex assembly. For example, in the hinge region, e.g., G24C G27C, Cys-substituted Hcf106 lost its ability to assemble into a 700 kDa receptor complex as analyzed by BN-PAGE, suggesting important contacts have been disrupted. This is in agreement with observations in *E. coli* TatB hinge where the Gly/Pro residues have been identified as essential for efficient substrate export (Barrett et al., 2003). There are at least three possible explanations for the absence of the 700 kDa complex when cysteines are substituted into the hinge of Hcf106. The first is that the substitution with Cys at those residues inhibited Hcf106-Hcf106 or Hcf106-cpTatC interactions, resulting in the absence of the 700 kDa complex. Second, Cys substitutions at those residues did not inhibit assembly *per se* but impacted the stability of the 700 kDa complex resulting in a receptor complex to be unable to withstand digitonin solubilization. Third, these substitutions resulted in decreased integration and therefore decreased participation in the 700 kDa complex. The third explanation is unlikely because Cys substitutions in the hinge region did not appear to inhibit integration and resistance to alkaline extraction of Hcf106. However, the data do suggest that the N terminus around residues G8 and E11 and the hinge region around residues G24 and G27 are of great structural significance to the assembly or stability of the interaction between Hcf106 and cpTatC. Glycine often introduces more flexibility to protein structure, and so in replacing glycine with cysteine, the overall flexibility of the hinge region might be changed to interfere with an interaction with cpTatC. In addition to glycine, residue charge has been known to play an important role in membrane protein stability, possibly by forming a salt bridge and oligomeric structure solubilization (Wimley et al., 1996). For example, Tha4 contains a glutamate (E10) in the transmembrane region that has been shown to be critical for Tha4 function, possibly by stabilization of oligomers through salt bridge formation (Dabney-Smith et al., 2003). However, whether E11 in Hcf106 has a similar role is unclear. Other Hcf106 residues, such as E41, Q43, and P44 may also affect interaction with cpTatC as shown by a decrease in the presence of those Hcf106 Cys variants in the 700 kDa complex. By replacing residues with Cys, Hcf106 contacts with cpTatC would be altered. Further studies are needed to clarify which region of cpTatC interacts with the N-terminus and hinge region of Hcf106. Currently, due to the existence of endogenous Hcf106, we are unable to determine whether these cysteine variants affect receptor complex functionality.

Truncated Hcf1061-107 can assemble with endogenous cpTatC indicating that the C-tail is less important as compared with the TMD and APH for receptor complex formation. This is also in agreement with the observation in *E.coli* that significant transport was observed when 70 residues were removed from C-terminus of TatB (Lee et al., 2002). We designed these experiments to maximize the observable amount of the 700 kDa receptor complex relative to homo-oligomeric Hcf106 complexes by adjusting the detergent:chlorophyll ratio. Based on our earlier BN-PAGE data, the homo-oligomeric forms should not appear on the gel. This allows us to interpret the lower bands seen when the truncated Hcf106 is incorporated into the receptor complex. As the amount of *in vitro* translated pH]Hcf1061-107 increases, more of the truncated protein assembled into a complex of lower molecular weight with cpTatC, indicating that Hcf1061-107 has the ability to compete with endogenous Hcf106 and as the concentration of Hcf1061-107 increases, more cpTatC assembled with the truncated Hcf106. However, based on the immunoblotting data, the proportion of full length Hcf106 in a 700 kDa complex is less. The apparent ~600 kDa and ~500 kDa bands, together with the double Cys cross-linking data, strongly suggest that integrated recombinant Hcf106 may form oligomers in a separate pool and may compete with the endogenous Hcf106 for association with cpTatC. If the receptor complex contains eight copies of both Hcf106 and cpTatC as predicted (Mori et al., 2001; Celedon and Cline, 2012), the ~600 kDa complexes may indicate approximately four endogenous Hcf106 were replaced with the truncated version while the ~500 kDa complexes indicates most, if not all, of the endogenous Hcf106 were replaced by truncated Hcf106.

Hcf106 shares structural similarity with Tha4. Both biochemical labeling (Aldridge et al., 2012) of Tha4 as well as solution NMR and computer simulation modeling (Rodriguez et al., 2013) of *E. coli* homolog TatA show that the TMD is tilted and the N-terminal APH is partially embedded in the membrane, rather than the TMD inserted vertically (e.g., TMD parallel to the bilayer normal) and the APH laying on the surface of membrane (e.g., perpendicular to the bilayer normal). The cross-linking data of Hcf106 presented here also suggest a similar topology for Hcf106. We saw limited interactions between residues L28C to R40C (N-proximal region of the APH), which could be explained by being in the low dielectric environment of the hydrophobic core of the membrane.

In summary, to systematically study Hcf106, a critical component in the chloroplast Tat system, a recombinant library of single Cys-substitutions from the N-terminus to the end of APH was generated. Here we demonstrate that exogenous, recombinant Hcf106 was able to insert into thylakoid and participates directly in the cpTat receptor complex, likely by replacing endogenous, Cys-less versions of Hcf106. This library not only helped clarify Hcf106 self-contacts but also demonstrated that exogenously integrated Hcf106 interacts with the precursor signal peptide via the Hcf106 APH and that the TMD was identified to form close contacts with cpTatC TM5. These interaction data further confirm the capability of *in vitro* integrated Hcf106 to function in the thylakoid membrane system and demonstrates a new tool to evaluate the organization of cpTat complexes.

## Acknowledgements

This work was supported by the Department of Energy: Office of Science Basic Energy Science (Award DE-SC0014441 to CDS). The authors wish to thank Ken Cline (University of Florida) for the cpTatC clones and members of the Dabney-Smith lab for critical reading of the manuscript.

## Author Contributions

QM and CDS designed the research; QM and KF performed research; QM, KF, CPN, and CDS analyzed data; QM, CPN, and CDS wrote the manuscript.

